# A jumbo cyanophage encodes the most complete ribosomal protein set in the known virosphere

**DOI:** 10.1101/2025.09.23.678156

**Authors:** Isaac Meza-Padilla, Sarit Avrani, Kirsten M. Müller, Jozef I. Nissimov

## Abstract

Viruses encode ribosomal proteins (RPs), but their genomes usually harbor a single RP-coding gene. Here, we reannotate the genome of the jumbo cyanophage PhiMa05 and show that it encodes six RPs, an RP acetyltransferase, and a ribosome biogenesis protein. Evolutionary analyses suggest that these viral RP-coding genes may have been horizontally transferred to certain members of the *Vampirovibrionia*, a non-photosynthetic basal lineage of *Cyanobacteriota*, via integration of the viral genome.

## Main

The ultimate divide between viruses and cells has been proposed to be the absence and presence of ribosomal genes, respectively^1^. This view was recently challenged by the discovery that viral genomes (including jumbo phages) occasionally encode ribosomal proteins (RPs)^2-8^. However, viruses usually harbor a single RP-coding gene^2-8^. Even giant Tupanviruses, possessing the most complete translational apparatus of the known virosphere, lack RPs altogether^9^. Thus far, to our knowledge, the viral RP-coding record has been held by eight uncultivated viruses encoding two RPs each^4^. Collectively, these intriguing findings prompt the following question: are there viruses in which selection has particularly favoured the acquisition of RP-coding genes?

We have recently identified the proteome of the cyanophage PhiMa05 while mining genomic data. PhiMa05 is a lytic jumbo phage that contains a ∼274 kb linear double-stranded DNA genome with 256 open reading frames (ORFs) and was isolated from an aeration basin of the wastewater treatment plant in a hospital in Thailand^10^. The virus infects seven toxic *Microcystis* strains isolated from the eutrophic, fresh-brackish-saline water Songkhla Lake^10,11^. Accordingly, whole-genome proteomic^12^, major capsid protein^10^, and portal protein (**Fig. S1**) phylogenies place PhiMa05 among freshwater cyanophages. In the study by Naknaen et al.^10^ examining the PhiMa05 genome, 54 ORFs were functionally annotated, however none were identified to be RPs (**Table S1**).

Our genome reannotation pipeline uncovered six RPs in the PhiMa05 genome, namely, bS1 (ORF 12), bL21 (ORF 13), bL27 (ORF 14), bL33 (ORF 35), uL11 (ORF 38) and uL1 (ORF 39), as well as a ribosome biogenesis GTP-binding YihA/YsxC protein (ORF 108), and an RP S18-alanine N-acetyltransferase (ORF 242; **Fig. 1, Table S1**). Specifically, NCBI Protein database mining rescued five RPs (bS1, bL21, bL27, uL11 and uL1); while BLASTP, InterProScan and Foldseek queries (**Table S2**) confirmed those five and revealed three more: bL33, the RP S18-alanine N-acetyltransferase, and the ribosome biogenesis protein (previously annotated only as a ‘GTP-binding protein’). It is worth noting that, except for uL11, the RPs encoded by PhiMa05 are located moderately close to one another in a prokaryotic ribosome (**Fig. 1**i). The viral genome also contains a few tRNA genes (tRNA-Trp, EMOOHJMP_00036; tRNA-Asn, EMOOHJMP_00158; and tRNA-Arg, EMOOHJMP_00211). To the best of our knowledge, this makes PhiMa05 the first cultivated cyanophage reported to encode RPs, as well as the virus with the most complete RP-coding set of the known virosphere.

**Fig. 1.**
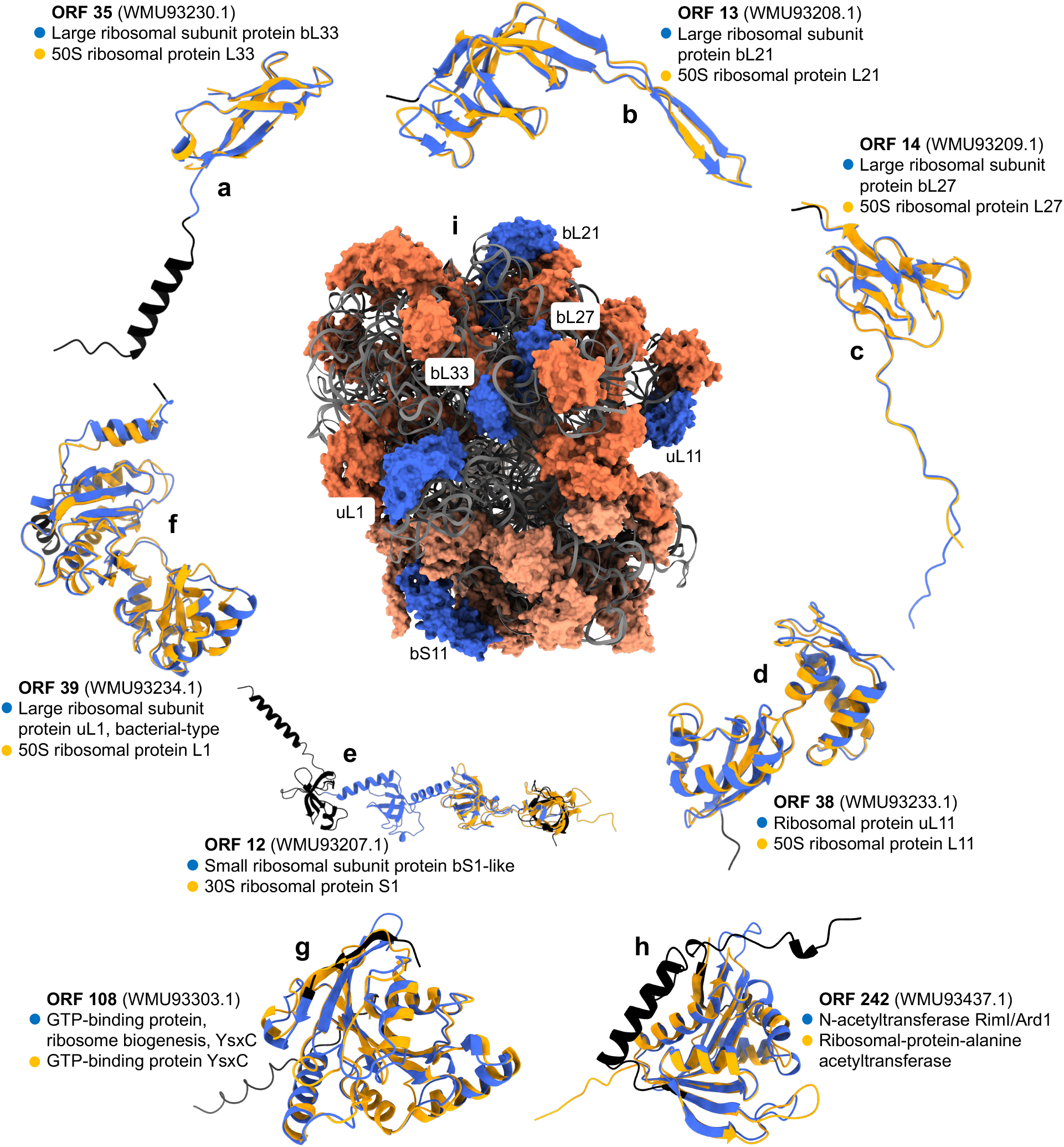
Ribosomal proteins (RPs) encoded in the genome of PhiMa05 and their locations in a bacterial ribosome. (a) bL33, (b) bL21, (c) bL27, (d) uL11, (e) bS1, (f) uL1, (g) ribosome biogenesis GTP-binding YihA/YsxC protein, and (h) RP S18-alanine N-acetyltransferase. (i) Shows a bacterial ribosome (PDB ID: 6BU8; from *Escherichia coli*) with the RPs encoded in the PhiMa05 genome colored in blue, the remaining large subunit RPs in coral, the remaining small subunit RPs in light salmon, and the RNAs in gray. For (a-h), the structural models of PhiMa05 proteins are colored in black, except for the regions corresponding to their InterPro families, which are colored in royal blue. The top Foldseek PDB hits are colored in orange and structurally aligned to the PhiMa05 structural models. The InterPro family names of the PhiMa05 proteins are displayed next to closed blue circles, while the names of their top Foldseek PDB hits are listed next to closed orange circles. Open reading frame (ORF) numbers of PhiMa05 proteins are highlighted in bold and their GenBank accession numbers are indicated in parentheses. Note that the different panels are not to scale. See **Table S1** for BLASTP *E*-values; **Table S2** for Foldseek *E*-values and InterPro family IDs; and **Fig. S2** for the predicted local-distance difference test scores of the structural models.

This fascinating discovery raises questions regarding the evolutionary trajectory of the PhiMa05 RP-coding genes. In all of our phylogenetic analyses, the RPs from PhiMa05 are strongly supported as being closely related to *Vampirovibrio* RPs (**Fig. 2**a-h). This suggests that the PhiMa05 RP-coding genes may have been acquired by an ancestor of the virus from a *Vampirovibrio*-like host. For context, *Vampirovibrio* is a genus of predatory cyanobacteria that belong to a basal non-photosynthetic class of the phylum *Cyanobacteriota*, the *Vampirovibrionia* (previously *Melainabacteria*)^13^. Interestingly, the comparative genomic analyses implemented here reveal that there is a highly conserved PhiMa05-like prophage in the genomes of *Vampirovibrio chlorellavorus* strains Vc_AZ_1 (GenBank accession number: GCA_003149375.1) and Vc_AZ_2 (**Fig. 2**i; GCA_003149345.1), which were isolated from outdoor experimental algal cultivation ponds in Arizona^14^. *Vampirovibrio chlorellavorus* is an obligate predator of the green microalga *Chlorella*^13,14^. Except for bL33 in the Vc_AZ_2 prophage, the rest of the PhiMa05 RP-coding genes are highly conserved in both prophages. Further, it is critical to note that uL1, uL11, bL21, bL27, the ribosome biogenesis GTP-binding YihA/YsxC, and the RP S18-alanine N-acetyltransferase of the PhiMa05-like prophages are the only copies of those proteins that the Vc_AZ_1 and Vc_AZ_2 host genomes possess. We hypothesize that these two cellular organisms may be totally dependent on the PhiMa05-like prophage for protein synthesis, and hence life itself.

**Fig. 2.**
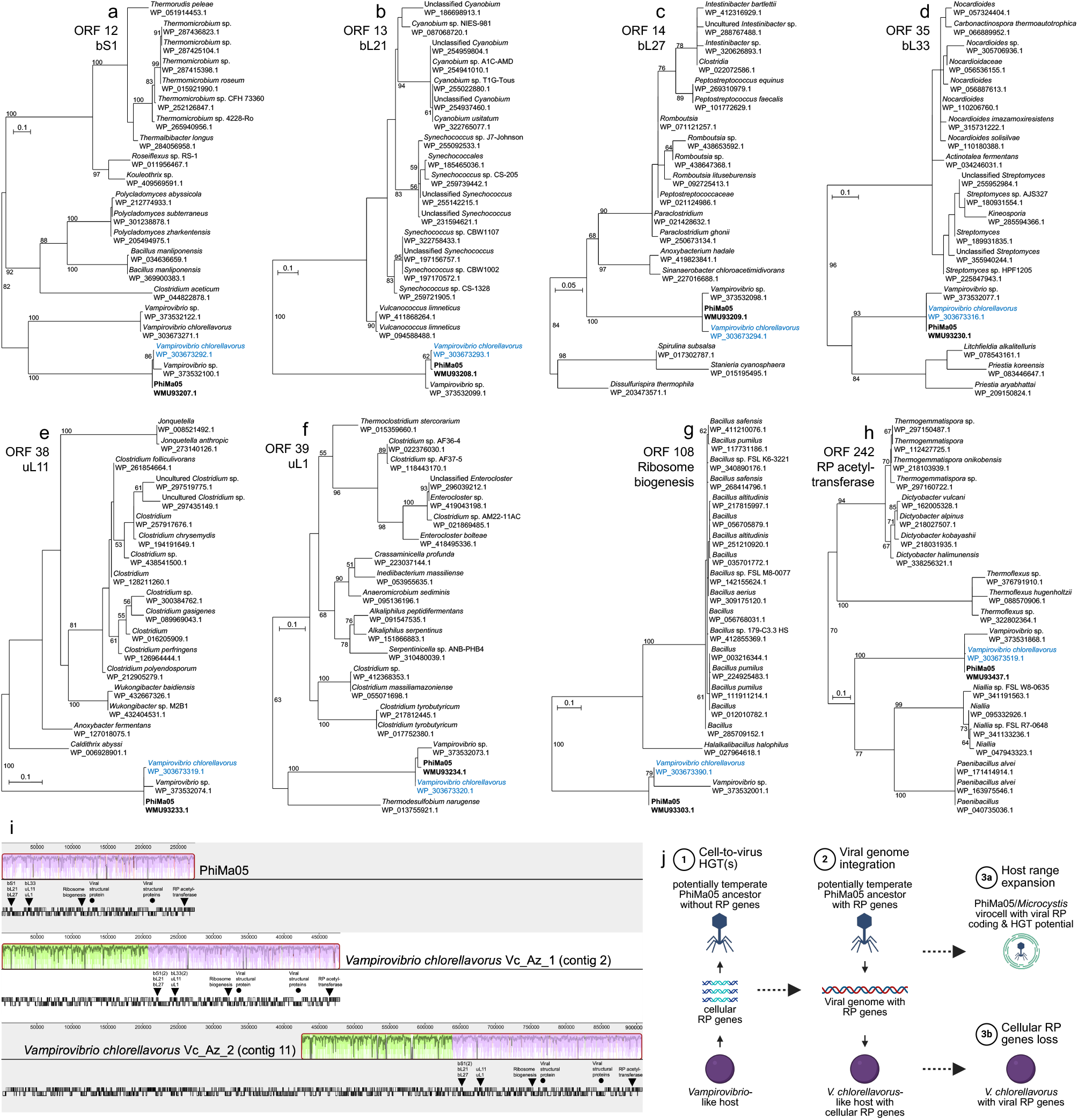
Maximum likelihood phylogenetic trees of PhiMa05 ribosomal proteins (RPs), synteny comparison between the PhiMa05 genome and *Vampirovibrio chlorellavorus* prophages, and evolutionary model. (a) bS1, (b) bL21, (c) bL27, (d) bL33, (e) uL11, (f) uL1, (g) ribosome biogenesis GTP-binding YihA/YsxC protein, and (h) RP S18-alanine N-acetyltransferase. The PhiMa05 open reading frame (ORF) numbers and protein names are indicated next to the trees. Support values > 50 (out of 100 bootstrap replicates) are shown at the nodes. The scale bars indicate the number of amino acid substitutions per site. RefSeq accession numbers are included below the species names. PhiMa05 proteins are highlighted in bold, and *V. chlorellavorus* Vc_AZ_1 PhiMa05-like prophage proteins are colored in blue. (i) Synteny analysis between PhiMa05 (top), *V. chlorellavorus* Vc_AZ_1 (middle) and *V. chlorellavorus* Vc_AZ_2 genomes (bottom). The locally colinear blocks (LCBs) corresponding to the PhiMa05 genome are colored in mauve, while the remaining syntenic regions between the Vc_AZ_1 and Vc_AZ_2 contigs are colored in green. The plots within the LCBs display sequence similarity values as a range with the mean value darkened. The coding sequences are displayed as black and white bars below each genome; PhiMa05 RPs and structural proteins (including tail, adaptor, capsid and portal proteins) are indicated using closed black triangles and circles, respectively. (j) Evolutionary model for the causes and consequences of PhiMa05 RP-coding genes (created in BioRender). See text for a more detailed description.

Based on the results presented here, we propose the following evolutionary scenario (**Fig. 2**j). First, a potentially temperate ancestor of PhiMa05 horizontally acquired RP-coding genes from a *Vampirovibrio*-like host. Second, the PhiMa05 ancestor integrated its genome in a *V. chlorellavorus* Vc_AZ_1/2-like host that probably carried its own cellular set of RP-coding genes. Third, the Vc_AZ_1/2-like host lost the cellular copies of the genes coding for uL1, uL11, bL21, bL27, the ribosome biogenesis, and the RP acetyltransferase proteins. As a result, the only genes for four RPs, a ribosome biogenesis protein, and an RP acetyltransferase encoded in the genomes of *V. chlorellavorus* Vc_AZ_1 and Vc_AZ_2 are of viral (prophage) origin. Whether these prophages are currently inducible remains unclear and should be investigated in future studies. Over the course of evolution, the PhiMa05 ancestor likely experienced an inter-class host range expansion or shift, leading to the current *Microcystis*-infecting PhiMa05 containing RP-coding genes. Given that jumbo phages have wider host ranges than smaller phages^15^, the close association of *Vampirovibrionales* strains with toxic *Microcystis* blooms^16^, and the existence of *Microcystis* phages with exceedingly broad host ranges^17^, such host expansion or shift would indeed be possible. Overall, our findings suggest that selection has particularly favoured the acquisition of RP-coding genes in certain viruses and that, in some cases, such viral genes have been transferred to cells, likely becoming essential for the survival of the host.

## Methods

Whole-genome functional reannotation of PhiMa05 (GenBank accession number: MW495066.1) was conducted using a combination of NCBI Protein database mining, RefSeq BLASTP searches and, when ribosomal proteins were detected, InterProScan^18^ runs, and Foldseek^19^ queries against experimentally determined structures in the PDB in order to confirm their annotations using multiple tools. Unless otherwise noted, all tools were executed with default parameters. BLASTP hits with *E*-values < 0.001 were considered significant, as were Foldseek hits with *E*-values < 0.001 and true-positive probabilities of 1^20,21^. Structural modelling for PhiMa05 RPs was conducted using AlphaFold 3^22^ (AF3), with the exception of bS1. The overall fold of AlphaFold models with predicted template modelling (pTM) scores > 0.5 is usually considered similar to that of the true structure^21^. Since AF3 failed to produce a satisfactory model for bS1 (pTM < 0.5), D-I-TASSER^23^ was used instead. To ensure optimal modelling accuracy, the following advanced options were activated in D-I-TASSER: using large IMG/JGI metagenomic database to build MSA, predict protein function based on structure model, using AlphaFold2 distances, and using domain partition and assembly module. As with AF3 pTM, an estimated template modelling (eTM) score > 0.5 has been used as threshold to denote correctly folded D-I-TASSER models^23^. Prior to Foldseek searches, the predicted local-distance difference test scores of the AF3 models were visually inspected in the native environment of the AlphaFold server and can be found in **Fig. S2**. The structural alignments generated by Foldseek were visualized using UCSF ChimeraX 1.10^24^.

For phylogenetic analyses, multiple sequence alignments of the top 20 RefSeq BLASTP hits of PhiMa05 RPs were computed using Muscle 5.3^25^ and visualized in SeaView^26^. Blocks of conserved sites were selected via Gblocks^27^, and maximum likelihood phylogenies were reconstructed using PhyML 3.1^28^ with the LG amino acid substitution model^29^ and 100 bootstrap replicates. In the case of the portal protein alignment, the use of Gblocks with default parameters yielded an insufficient number of positions, resulting in a tree with low bootstrap support for some branches. Therefore, Gblocks was executed with the following parameters to increase the robustness of the portal protein tree: allow smaller final blocks, allow gap positions within the final blocks, and allow less strict flanking positions. Finally, a multiple genome alignment between PhiMa05, *Vampirovibrio chlorellavorus* Vc_AZ_1 (contig QFWH01000002.1) and *V. chlorellavorus* Vc_AZ_2 (contig QFWI01000011.1) was performed using progressiveMauve^30^. Since the *V. chlorellavorus* Vc_AZ_2 genome was not annotated in NCBI, Prokka 1.14.5^31^ was used to annotate it before alignment.

## Acknowledgements

This work was funded by a Secretaría de Ciencia, Humanidades, Tecnología e Innovación (SECIHTI) Becas de Posgrado para Maestrías y Doctorados en Ciencias y Humanidades en el Extranjero scholarship awarded to I.M.P.; as well as Natural Sciences and Engineering Research Council of Canada (NSERC) Discovery Grants (2022-03350 and 2022-00329), and a Phycological Society of America (PSA) Norma J. Lang Early Career Researcher Fellowship awarded to J.I.N.

## Author Contributions

I.M.P. conceived the idea, conducted the bioinformatics pipeline, analyzed the data and wrote the manuscript. S.A., K.M.M. and J.I.N. provided valuable advice and feedback on the manuscript draft. I.M.P and J.I.N. acquired funding. All authors revised and edited the manuscript.

