## Supplementary Material for "A jumbo cyanophage encodes the most complete ribosomal protein set in the known virosphere"

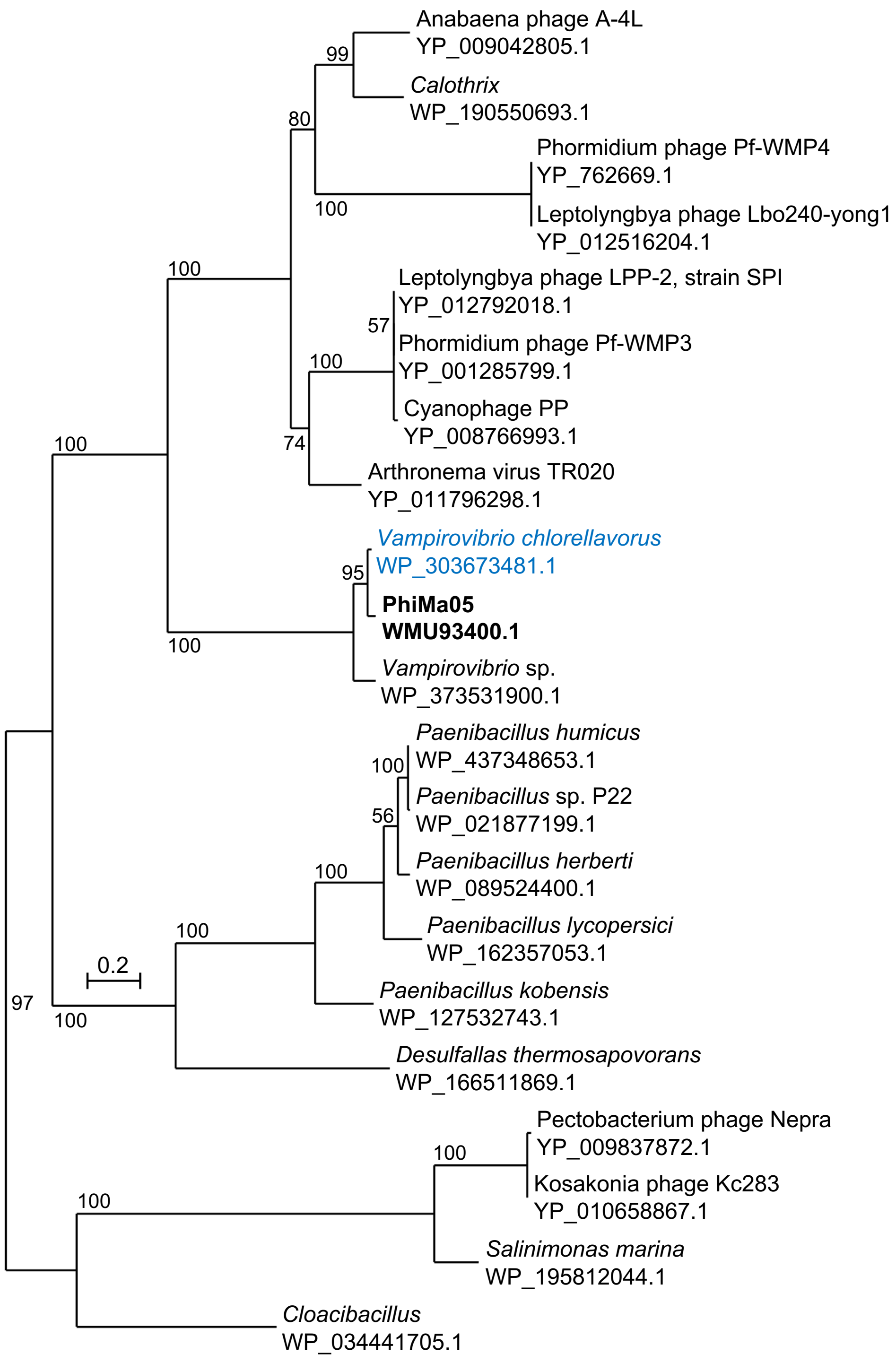


**Fig. S1**. Maximum likelihood phylogenetic tree of PhiMa05 portal protein. Support values > 50 (out of 100 bootstrap replicates) are shown at the nodes. The scale bar indicates the number of amino acid substitutions per site. RefSeq accession numbers are included below the species names. PhiMa05 portal protein is highlighted in bold, and *Vampirovibrio chlorellavorus* Vc_AZ_1 PhiMa05-like prophage portal protein is colored in blue.

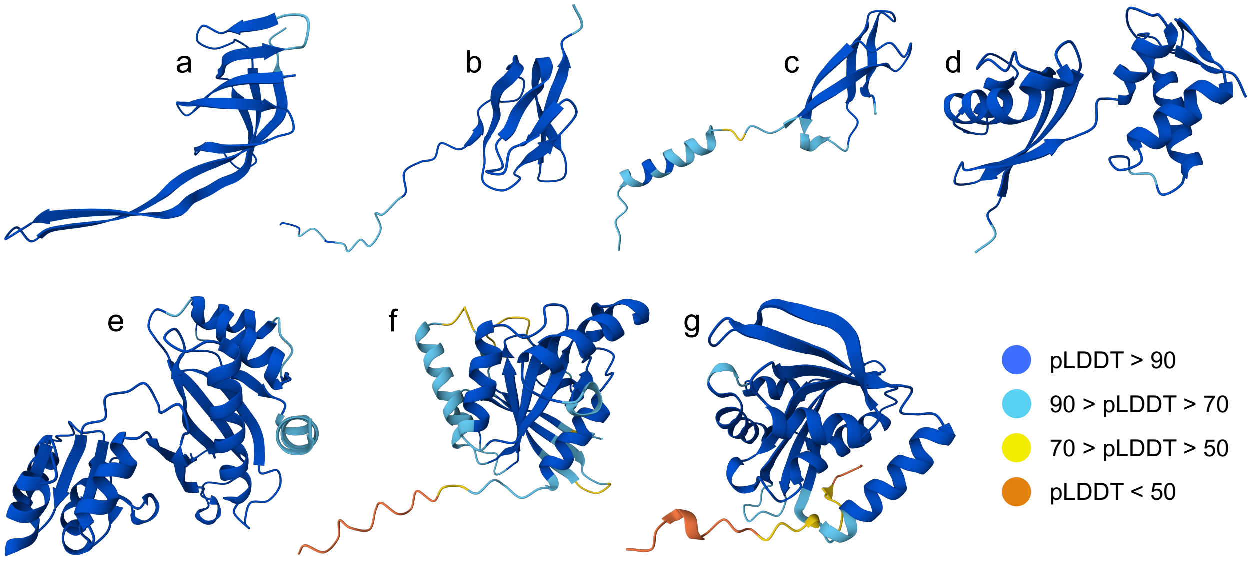


**Fig. S2**. Predicted local-distance difference test (pLDDT) scores of the AlphaFold 3 structural models of PhiMa05 ribosomal proteins (RPs). (a) bL21, (b) bL27, (c) bL33, (d) uL11, (e) uL1, (f) ribosome biogenesis GTP-binding YihA/YsxC protein, and (g) RP S18-alanine N-acetyltransferase. The different panels are not to scale. bS1 is not shown because D-I-TASSER models do not include pLDDT scores. However, the estimated template modelling score of the bS1 structural model can be found in **Table S2**.

**Table S1**. Annotations and homologs of PhiMa05 open reading frames. Ribosomal proteins are underlined.

| ORF | GenBank accession | Naknaen et al.[10] annotation[a] (2021 paper) | GenBank annotation (2023 revision) | RefSeq homolog[b] (this study) | Organism (this study) | BLASTP *E*-value (this study) |
|---|---|---|---|---|---|---|
| 1 | WMU93196.1 | 4-hydroxy-3-methylbut-2-enyl diphosphate synthase | 4-hydroxy-3-methylbut-2-en-1-yl diphosphate synthase | Flavodoxin-dependent (E)-4-hydroxy-3-methylbut-2-enyl-diphosphate synthase | Unclassified *Clostridium* | 1e-129 |
| 2 | WMU93197.1 | N.a. | HP | 2 HPs | *Vampirovibrio* spp. | ≤ 2e-45 |
| 3 | WMU93198.1 | N.a. | HP | Radical SAM protein | *V. chlorellavorus* | 0 |
| 4 | WMU93199.1 | N.a. | Cyclic dehypoxanthine futalosine synthase | Cyclic dehypoxanthinyl futalosine synthase | *V. chlorellavorus* | 0 |
| 5 | WMU93200.1 | N.a. | HP | 2 HPs | *Vampirovibrio* spp. | ≤ 5e-96 |
| 6 | WMU93201.1 | N.a. | HP | 2 HPs | *Vampirovibrio* spp. | ≤ 1e-38 |
| 7 | WMU93202.1 | Tryptophanyl-tRNA synthetase | Tryptophan--tRNA ligase | Tryptophan--tRNA ligase | *V. chlorellavorus* | 0 |
| 8 | WMU93203.1 | N.a. | HP | 2 HPs | *Vampirovibrio* spp. | ≤ 2e-104 |
| 9 | WMU93204.1 | N.a. | HP | Prepilin peptidase | *V. chlorellavorus* | 0 |
| 10 | WMU93205.1 | Putative RNA polymerase sigma subunit | RNA polymerase sigma factor | Sigma-70 family RNA polymerase sigma factor | *V. chlorellavorus* | 0 |
| 11 | WMU93206.1 | N.a. | 4-hydroxy-3-methylbut-2-enyl diphosphate reductase | 4-hydroxy-3-methylbut-2-enyl diphosphate reductase | *V. chlorellavorus* | 0 |
| 12 | WMU93207.1 | N.a. | 30S ribosomal protein S1 | 30S ribosomal protein S1 | *V. chlorellavorus* | 0 |
| 13 | WMU93208.1 | N.a. | 50S ribosomal protein L21 | 50S ribosomal protein L21 | *V. chlorellavorus* | 2e-69 |
| 14 | WMU93209.1 | N.a. | 50S ribosomal protein L27 | 50S ribosomal protein L27 | *V. chlorellavorus* | 3e-53 |
| 15 | WMU93210.1 | N.a. | HP | 2 HPs | *Vampirovibrio* spp. | ≤ 4e-13 |
| 16 | WMU93211.1 | N.a. | Holliday junction ATP-dependent DNA helicase | Holliday junction branch migration protein RuvA | *V. chlorellavorus* | 5e-142 |
| 17 | WMU93212.1 | N.a. | 2-C-methyl-D-erythritol 4-phosphate cytidylyltransferase | 2-C-methyl-D-erythritol 4-phosphate cytidylyltransferase | *V. chlorellavorus* | 7e-103 |
| 18 | WMU93213.1 | N.a. | Thiamine-monophosphate kinase | Thiamine-phosphate kinase | *V. chlorellavorus* | 0 |
| 19 | WMU93214.1 | N.a. | HP | 2 HPs | *Vampirovibrio* spp. | ≤ 2e-07 |
| 20 | WMU93215.1 | Aerotaxis sensor receptor protein | HP | Globin-coupled sensor protein | *V. chlorellavorus* | 0 |
| 21 | WMU93216.1 | N.a. | HP | Type II secretion system F family protein | *V. chlorellavorus* | 0 |
| 22 | WMU93217.1 | N.a. | HP | Type II secretion system F family protein | *V. chlorellavorus* | 0 |
| 23 | WMU93218.1 | CpaF Flp pilus assembly protein, ATPase CpaF | HP | CpaF family protein | *V. chlorellavorus* | 0 |
| 24 | WMU93219.1 | N.a. | Protein-glutamate methylesterase/ protein-glutamine glutaminase | AAA family ATPase | *V. chlorellavorus* | 0 |
| 25 | WMU93220.1 | N.a. | Type 3 secretion system secretin | Type II and III secretion system protein family protein | *V. chlorellavorus* | 0 |
| 26 | WMU93221.1 | N.a. | HP | Flp pilus assembly protein CpaB | *V. chlorellavorus* | 0 |
| 27 | WMU93222.1 | N.a. | HP | TadE family protein | *V. chlorellavorus* | 2e-83 |
| 28 | WMU93223.1 | N.a. | HP | 3 HPs | *Vampirovibrio* spp. | ≤ 7e-09 |

| | | | | | | |
|---|---|---|---|---|---|---|
| 29 | WMU93224.1 | Thioredoxin reductase gp344 | Thioredoxin reductase | Thioredoxin-disulfide reductase | *V. chlorellavorus* | |
| 30 | WMU93225.1 | N.a. | HP | 2 HPs | *Vampirovibrio* spp. | ≤ 4e-13 |
| 31 | WMU93226.1 | N.a. | 3'3'-cGAMP-specific phosphodiesterase 3 | HD-GYP domain-containing protein | *V. chlorellavorus* | 0 |
| 32 | WMU93227.1 | N.a. | HP | DivIVA domain-containing protein | *Zarconia navalis* | 2e-26 |
| 33 | WMU93228.1 | N.a. | HP | 2 HPs | *Vampirovibrio* spp. | ≤ 3e-52 |
| 34 | WMU93229.1 | CoaD Phosphopantetheine adenylyltransferase | Phosphopantetheine adenylyltransferase | Pantetheine-phosphate adenylyltransferase | *V. chlorellavorus* | 8e-114 |
| 35 | WMU93230.1 | N.a. | HP | 50S ribosomal protein L33 | *V. chlorellavorus* | 2e-31 |
| 36 | WMU93231.1 | N.a. | Protein translocase subunit | Preprotein translocase subunit SecE | *V. chlorellavorus* | 2e-35 |
| 37 | WMU93232.1 | Transcription antitermination protein | Transcription termination/antitermination protein | Transcription termination/antitermination protein NusG | *Vampirovibrio* sp. | 9e-151 |
| 38 | WMU93233.1 | N.a. | 50S ribosomal protein L11 | 50S ribosomal protein L11 | *V. chlorellavorus* | 9e-98 |
| 39 | WMU93234.1 | N.a. | 50S ribosomal protein L1 | 50S ribosomal protein L1 | *V. chlorellavorus* | 2e-169 |
| 40 | WMU93235.1 | AhpC Peroxiredoxin | Selenocysteine-containing peroxiredoxin | Peroxiredoxin | *V. chlorellavorus* | 2e-126 |
| 41 | WMU93236.1 | N.a. | Thiol-disulfide oxidoreductase | Peroxiredoxin family protein | *V. chlorellavorus* | 5e-68 |
| 42 | WMU93237.1 | N.a. | 2-dehydro-3-deoxygluconokinase | Sugar kinase | *V. chlorellavorus* | 0 |
| 43 | WMU93238.1 | N.a. | Dihydroxy-acid dehydratase | Dihydroxy-acid dehydratase | *V. chlorellavorus* | 0 |
| 44 | WMU93239.1 | N.a. | KHG/KDPG aldolase | Bifunctional 4-hydroxy-2-oxoglutarate aldolase/2-dehydro-3-deoxy-phosphogluconate aldolase | *V. chlorellavorus* | 5e-137 |
| 45 | WMU93240.1 | N.a. | Kynurenine formamidase | Cyclase family protein | *V. chlorellavorus* | 4e-153 |
| 46 | WMU93241.1 | N.a. | HP | KGG domain-containing protein | *V. chlorellavorus* | 8e-38 |
| 47 | WMU93242.1 | N.a. | Epoxyqueuosine reductase | tRNA epoxyqueuosine(34) reductase QueG | *V. chlorellavorus* | 0 |
| 48 | WMU93243.1 | N.a. | Glucose-6-phosphate isomerase | Glucose-6-phosphate isomerase | *V. chlorellavorus* | 0 |
| 49 | WMU93244.1 | Ferredoxin/ferredoxin--NADP reductase | Ferredoxin | 4Fe-4S dicluster domain-containing protein | *V. chlorellavorus* | 1e-52 |
| 50 | WMU93245.1 | N.a. | HP | 2 HPs | *Vampirovibrio* spp. | ≤ 2e-25 |
| 51 | WMU93246.1 | Sensor domain-containing diguanylate cyclase | HP | GGDEF domain-containing protein | *V. chlorellavorus* | 0 |
| 52 | WMU93247.1 | N.a. | Cysteine desulfurase | Aminotransferase class V-fold PLP-dependent enzyme | *V. chlorellavorus* | 0 |
| 53 | WMU93248.1 | N.a. | HP | Cysteine dioxygenase | *V. chlorellavorus* | 2e-106 |
| 54 | WMU93249.1 | N.a. | Heme chaperone | Radical SAM family heme chaperone HemW | *V. chlorellavorus* | 0 |
| 55 | WMU93250.1 | N.a. | HP | HP | *V. chlorellavorus* | 1e-34 |
| 56 | WMU93251.1 | N.a. | HP | 2 HPs | *Vampirovibrio* spp. | ≤ 1e-26 |
| 57 | WMU93252.1 | Isocitrate dehydrogenase, partial | Isocitrate dehydrogenase [NADP] | Isocitrate dehydrogenase (NADP(+)) | *V. chlorellavorus* | 0 |
| 58 | WMU93253.1 | N.a. | Aconitate hydratase A | Aconitate hydratase | *V. chlorellavorus* | 0 |

| | | | | | | |
|---|---|---|---|---|---|---|
| 59 | WMU93254.1 | Ribonucleoside-diphosphate reductase | Vitamin B12-dependent ribonucleoside-diphosphate reductase | Adenosylcobalamin-dependent ribonucleoside-diphosphate reductase | *V. chlorellavorus* | 0 |
| 60 | WMU93255.1 | N.a. | D-inositol-3-phosphate glycosyltransferase | Glycosyltransferase family 4 protein | *V. chlorellavorus* | 0 |
| 61 | WMU93256.1 | RNA polymerase sigma factor | RNA polymerase sigma factor | Sigma-70 family RNA polymerase sigma factor | *V. chlorellavorus* | 0 |
| 62 | WMU93257.1 | N.a. | Photosystem I assembly protein | Tetratricopeptide repeat protein | *V. chlorellavorus* | 0 |
| 63 | WMU93258.1 | N.a. | Ribonuclease Y | HDOD domain-containing protein | *V. chlorellavorus* | 0 |
| 64 | WMU93259.1 | N.a. | HP | 2 HPs | *Vampirovibrio* spp. | ≤ 1e-89 |
| 65 | WMU93260.1 | N.a. | HP | Flagellar filament capping protein FliD | *V. chlorellavorus* | 0 |
| 66 | WMU93261.1 | N.a. | Putative amino acid permease | Aminoacid permease | *V. chlorellavorus* | 0 |
| 67 | WMU93262.1 | N.a. | HP | 2 HPs | *Vampirovibrio* spp. | ≤ 7e-27 |
| 68 | WMU93263.1 | N.a. | tRNA-specific 2-thiouridylase | tRNA 2-thiouridine(34) synthase MnmA | *V. chlorellavorus* | 0 |
| 69 | WMU93264.1 | N.a. | HP | Pentapeptide repeat-containing protein | *V. chlorellavorus* | 0 |
| 70 | WMU93265.1 | N.a. | HP | M2 family metallopeptidase | *V. chlorellavorus* | 0 |
| 71 | WMU93266.1 | N.a. | HP | 2 HPs | *Vampirovibrio* spp. | ≤ 2e-61 |
| 72 | WMU93267.1 | Bifunctional NAD-dependent-3-hydroxypropionate dehydrogenase | Putative oxidoreductase | SDR family oxidoreductase | *V. chlorellavorus* | 0 |
| 73 | WMU93268.1 | N.a. | HP | 2 HPs | *Vampirovibrio* spp. | ≤ 4e-78 |
| 74 | WMU93269.1 | N.a. | HP | 3 HPs | *Vampirovibrio* spp. | ≤ 1e-04 |
| 75 | WMU93270.1 | N.a. | HP | HP | *V. chlorellavorus* | 8e-15 |
| 76 | WMU93271.1 | N.a. | HP | DUF3228 family protein | *V. chlorellavorus* | 8e-106 |
| 77 | WMU93272.1 | DNA polymerase IV | DNA polymerase IV | DNA polymerase IV | *V. chlorellavorus* | 0 |
| 78 | WMU93273.1 | N.a. | HP | HP | *V. chlorellavorus* | 4e-23 |
| 79 | WMU93274.1 | N.a. | HP | 2 HPs | *Vampirovibrio* spp. | ≤ 2e-20 |
| 80 | WMU93275.1 | N.a. | HP | 2 HPs | *Vampirovibrio* spp. | ≤ 7e-54 |
| 81 | WMU93276.1 | PrsA Phosphoribosyl-pyrophosphate synthetase | Ribose-phosphate pyrophosphokinase | Ribose-phosphate diphosphokinase | *V. chlorellavorus* | 0 |
| 82 | WMU93277.1 | N.a. | HP | AtaL-like protein | *V. chlorellavorus* | 1e-102 |
| 83 | WMU93278.1 | N.a. | HP | HP | *V. chlorellavorus* | 1e-139 |
| 84 | WMU93279.1 | N.a. | Anthranilate synthase component 2 | Anthranilate synthase component II | *V. chlorellavorus* | 1e-119 |
| 85 | WMU93280.1 | N.a. | HP | HP | *V. chlorellavorus* | 2e-73 |
| 86 | WMU93281.1 | N.a. | HP | 2 HPs | *Vampirovibrio* spp. | ≤ 3e-54 |
| 87 | WMU93282.1 | N.a. | HP | 2 HPs | *Vampirovibrio* spp. | 0 |
| 88 | WMU93283.1 | N.a. | HP | ABC transporter permease | *V. chlorellavorus* | 0 |
| 89 | WMU93284.1 | N.a. | Glutathione transport system permease protein | ABC transporter permease | *V. chlorellavorus* | 0 |
| 90 | WMU93285.1 | N.a. | Acetyltransferase | GNAT family N-acetyltransferase | *V. chlorellavorus* | 1e-94 |
| 91 | WMU93286.1 | N.a. | Putative protein YccU | CoA-binding protein | *V. chlorellavorus* | 4e-85 |
| 92 | WMU93287.1 | N.a. | HP | HP | *V. chlorellavorus* | 9e-157 |

| | | | | | | |
|---|---|---|---|---|---|---|
| 93 | WMU93288.1 | 3-polyprenyl-4-hydroxybenzoate carboxy-lyase | Putative UbiX-like flavin prenyltransferase | UbiX family flavin prenyltransferase | *V. chlorellavorus* | 6e-158 |
| 94 | WMU93289.1 | N.a. | HP | 2 HPs | *Vampirovibrio* spp. | ≤ 1e-61 |
| 95 | WMU93290.1 | PcnB tRNA nucleotidyl-transferase/poly(A) polymerase | Poly(A) polymerase I | CCA tRNA nucleotidyltransferase | *Desulfobacula* sp. | 3e-42 |
| 96 | WMU93291.1 | N.a. | ATP-dependent dethiobiotin synthetase | AAA family ATPase | *V. chlorellavorus* | 1e-168 |
| 97 | WMU93292.1 | N.a. | Lipid II flippase | Murein biosynthesis integral membrane protein MurJ | *V. chlorellavorus* | 0 |
| 98 | WMU93293.1 | Proline dehydrogenase | Proline dehydrogenase 1 | Proline dehydrogenase family protein | *V. chlorellavorus* | 0 |
| 99 | WMU93294.1 | N.a. | Acyl-CoA dehydrogenase | Acyl-CoA dehydrogenase family protein | *V. chlorellavorus* | 0 |
| 100 | WMU93295.1 | N.a. | Chemotaxis protein | Motility protein A | *V. chlorellavorus* | 4e-180 |
| 101 | WMU93296.1 | N.a. | Outer membrane protein A | OmpA/MotB family protein | *V. chlorellavorus* | 3e-152 |
| 102 | WMU93297.1 | N.a. | HP | Flagellar basal body-associated FliL family protein | *V. chlorellavorus* | 4e-153 |
| 103 | WMU93298.1 | N.a. | Flagellar motor switch protein | Flagellar motor switch protein FliM | *V. chlorellavorus* | 0 |
| 104 | WMU93299.1 | N.a. | HP | Flagellar motor switch phosphatase FliY | *V. chlorellavorus* | 0 |
| 105 | WMU93300.1 | N.a. | Secreted effector protein | Pentapeptide repeat-containing protein | *V. chlorellavorus* | 7e-169 |
| 106 | WMU93301.1 | N.a. | HP | Metallothionein | *V. chlorellavorus* | 8e-45 |
| 107 | WMU93302.1 | Enoyl-[acyl-carrier-protein] reductase [NADH] | Enoyl-[acyl-carrier-protein] reductase [NADH] | Enoyl-ACP reductase FabI | *V. chlorellavorus* | 0 |
| 108 | WMU93303.1 | GTP-binding protein | Putative GTP-binding protein | Ribosome biogenesis GTP-binding protein YihA/YsxC | *V. chlorellavorus* | 9e-151 |
| 109 | WMU93304.1 | N.a. | HP | SH3 domain-containing protein | *Geomonas oryzisoli* | 8e-05 |
| 110 | WMU93305.1 | N.a. | HP | 2 HPs | *Vampirovibrio* spp. | ≤ 4e-21 |
| 111 | WMU93306.1 | N.a. | HP | 2 HPs | *Vampirovibrio* spp. | ≤ 2e-52 |
| 112 | WMU93307.1 | N.a. | HP | DUF366 family protein | *V. chlorellavorus* | 8e-138 |
| 113 | WMU93308.1 | Penicillin-insensitive trans-glycosylase & transpeptidase | Biosynthetic peptidoglycan transglycosylase | Transglycosylase domain-containing protein | *V. chlorellavorus* | 0 |
| 114 | WMU93309.1 | Glycine-tRNA ligase beta subunit | Glycine--tRNA ligase beta subunit | Glycine--tRNA ligase subunit beta | *V. chlorellavorus* | 0 |
| 115 | WMU93310.1 | N.a. | Lipid II isoglutaminyl synthase (glutamine-hydrolyzing) subunit | Type 1 glutamine amidotransferase | *V. chlorellavorus* | 7e-160 |
| 116 | WMU93311.1 | N.a. | HP | 2 HPs | *Vampirovibrio* spp. | ≤ 9e-22 |
| 117 | WMU93312.1 | N.a. | HP | 2 HPs | *Vampirovibrio* spp. | ≤ 3e-43 |
| 118 | WMU93313.1 | N.a. | HP | TIGR04552 family protein | *V. chlorellavorus* | 0 |
| 119 | WMU93314.1 | N.a. | Protein-export membrane protein | Protein translocase subunit SecF | *V. chlorellavorus* | 0 |
| 120 | WMU93315.1 | N.a. | Protein translocase subunit | Protein translocase subunit SecD | *V. chlorellavorus* | 0 |
| 121 | WMU93316.1 | Tail fibers protein | HP | Phosphodiester glycosidase family protein | *V. chlorellavorus* | 0 |
| 122 | WMU93317.1 | N.a. | HP | 2 HPs | *Vampirovibrio* spp. | ≤ 1e-95 |
| 123 | WMU93318.1 | N.a. | HP | 2 HPs | *Vampirovibrio spp.* | ≤ 7e-102 |
| 124 | WMU93319.1 | N.a. | Holo-[acyl-carrier-protein] synthase | Holo-ACP synthase | *V. chlorellavorus* | 5e-90 |

| | | | | | | |
|---|---|---|---|---|---|---|
| 125 | WMU93320.1 | N.a. | HP | 2 HPs | *Vampirovibrio* spp. | ≤ 3e-56 |
| 126 | WMU93321.1 | N.a. | HP | Serine/threonine-protein kinase | Unclassified *Streptosporangium* | 1e-12 |
| 127 | WMU93322.1 | N.a. | HP | 2 HPs | *Vampirovibrio* spp. | ≤ 1e-151 |
| 128 | WMU93323.1 | N.a. | HP | 2 HPs | *Vampirovibrio* spp. | ≤ 1e-35 |
| 129 | WMU93324.1 | N.a. | HP | AbrB/MazE/SpoVT family DNA-binding domain-containing protein | *V. chlorellavorus* | 1e-09 |
| 130 | WMU93325.1 | N.a. | HP | Type II toxin-antitoxin system PemK/MazF family toxin | *Ekhidna* sp. | 1e-15 |
| 131 | WMU93326.1 | N.a. | Segregation and condensation protein B | SMC-Scp complex subunit ScpB | *V. chlorellavorus* | 6e-155 |
| 132 | WMU93327.1 | N.a. | Segregation and condensation protein A | Segregation and condensation protein A | *V. chlorellavorus* | 0 |
| 133 | WMU93328.1 | DNA repair exonuclease SbcCD ATPase subunit | Chromosome partition protein | Chromosome segregation protein SMC | *V. chlorellavorus* | 0 |
| 134 | WMU93329.1 | N.a. | Putative zinc protease | M16 family metallopeptidase | *V. chlorellavorus* | 0 |
| 135 | WMU93330.1 | Putative Zn-dependent peptidase | Putative zinc protease | M16 family metallopeptidase | *V. chlorellavorus* | 0 |
| 136 | WMU93331.1 | N.a. | HP | S8 family serine peptidase | *V. chlorellavorus* | 0 |
| 137 | WMU93332.1 | N.a. | HP | None | | |
| 138 | WMU93333.1 | GIY-YIG nuclease superfamily protein | HP | GIY-YIG nuclease family protein | *V. chlorellavorus* | 6e-54 |
| 139 | WMU93334.1 | N.a. | Cardiolipin synthase A | Phospholipase D-like domain-containing protein | *V. chlorellavorus* | 0 |
| 140 | WMU93335.1 | N.a. | HP | 2 HPs | *Vampirovibrio* spp. | ≤ 1e-22 |
| 141 | WMU93336.1 | N.a. | HP | Type II secretion system protein | *Vampirovibrio* sp. | 1e-52 |
| 142 | WMU93337.1 | N.a. | HP | Type II secretion system protein | *Vampirovibrio* sp. | 1e-45 |
| 143 | WMU93338.1 | N.a. | UvrABC system protein A | Excinuclease ABC subunit UvrA | *V. chlorellavorus* | 0 |
| 144 | WMU93339.1 | N.a. | HP | 2 HPs | *Vampirovibrio* spp. | ≤ 1e-109 |
| 145 | WMU93340.1 | N.a. | HP | 2 HPs | *Vampirovibrio* spp. | ≤ 3e-88 |
| 146 | WMU93341.1 | MutT-like nucleotide pyrophosphohydrolase | RNA pyrophosphohydrolase | NUDIX domain-containing protein | *V. chlorellavorus* | 2e-100 |
| 147 | WMU93342.1 | N.a. | HP | 2 HPs | *Vampirovibrio* spp. | ≤ 1e-06 |
| 148 | WMU93343.1 | N.a. | HP | 2 HPs | *Vampirovibrio* spp. | ≤ 5e-69 |
| 149 | WMU93344.1 | N.a. | UDP-N-acetylglucosamine 2-epimerase | Non-hydrolyzing UDP-N-acetylglucosamine 2-epimerase | *V. chlorellavorus* | 0 |
| 150 | WMU93345.1 | N.a. | HP | TonB family protein | *V. chlorellavorus* | 1e-180 |
| 151 | WMU93346.1 | TopA topoisomerase IA | DNA topoisomerase 1 | Type I DNA topoisomerase | *V. chlorellavorus* | 0 |
| 152 | WMU93347.1 | N.a. | HP | S-layer homology domain-containing protein | *Vampirovibrio* sp. | 2e-19 |
| 153 | WMU93348.1 | N.a. | Photosystem II 12 kDa extrinsic protein | ComEA family DNA-binding protein | *V. chlorellavorus* | 6e-58 |
| 154 | WMU93349.1 | N.a. | HP | 2 HPs | *Vampirovibrio* spp. | ≤ 4e-82 |
| 155 | WMU93350.1 | N.a. | HP | YggT family protein | *V. chlorellavorus* | 4e-58 |

| | | | | | | |
|---|---|---|---|---|---|---|
| 156 | WMU93351.1 | N.a. | S-methyl-5'-thioadenosine phosphorylase | S-methyl-5'-thioadenosine phosphorylase | *V. chlorellavorus* | 0 |
| 157 | WMU93352.1 | N.a. | HP | HP | *V. chlorellavorus* | 0 |
| 158 | WMU93353.1 | N.a. | HP | 2 HPs | *Vampirovibrio* spp. | ≤ 6e-15 |
| 159 | WMU93354.1 | N.a. | Cobalt/magnesium transport protein | Magnesium/cobalt transporter CorA | *V. chlorellavorus* | 0 |
| 160 | WMU93355.1 | N.a. | HP | 2 HPs | *Vampirovibrio* spp. | ≤ 4e-53 |
| 161 | WMU93356.1 | N.a. | HP | DUF2145 domain-containing protein | *V. chlorellavorus* | 1e-164 |
| 162 | WMU93357.1 | N.a. | HP | 4 HPs | *Vampirovibrio* spp. | ≤ 3e-14 |
| 163 | WMU93358.1 | N.a. | HP | 2 HPs | *Vampirovibrio* spp. | ≤ 3e-37 |
| 164 | WMU93359.1 | N.a. | Mannosylglucosyl-3-phosphoglycerate phosphatase | Bifunctional metallophosphatase/5'-nucleotidase | *V. chlorellavorus* | 0 |
| 165 | WMU93360.1 | N.a. | Zinc uptake regulation protein | Fur family transcriptional regulator | *V. chlorellavorus* | 1e-94 |
| 166 | WMU93361.1 | N.a. | HP | RNA recognition motif domain-containing protein | *V. chlorellavorus* | 2e-114 |
| 167 | WMU93362.1 | N.a. | HP | 4 HPs | *Vampirovibrio* spp. | ≤ 4e-08 |
| 168 | WMU93363.1 | N.a. | HP | Glycosyl hydrolase family 8 | *V. chlorellavorus* | 0 |
| 169 | WMU93364.1 | N.a. | HP | Sugar transferase | *V. chlorellavorus* | 2e-142 |
| 170 | WMU93365.1 | N.a. | HP | HP | *V. chlorellavorus* | 6e-136 |
| 171 | WMU93366.1 | N.a. | HP | 204 HPs | Multiple species | ≤ 4e-06 |
| 172 | WMU93367.1 | N.a. | Methylthioribulose-1-phosphate dehydratase | Methylthioribulose 1-phosphate dehydratase | *V. chlorellavorus* | 1e-138 |
| 173 | WMU93368.1 | N.a. | Acireductone dioxygenase | 1,2-dihydroxy-3-keto-5-methylthiopentene dioxygenase | *V. chlorellavorus* | 4e-121 |
| 174 | WMU93369.1 | N.a. | Enolase-phosphatase E1 | Acireductone synthase | *V. chlorellavorus* | 4e-149 |
| 175 | WMU93370.1 | Aldehyde dehydrogenase B | Succinate-semialdehyde dehydrogenase [NADP(+)] | Aldehyde dehydrogenase family protein | *V. chlorellavorus* | 0 |
| 176 | WMU93371.1 | N.a. | HP | ABC transporter substrate-binding protein | *V. chlorellavorus* | 0 |
| 177 | WMU93372.1 | N.a. | HP | 2 HPs | *Vampirovibrio* spp. | ≤ 3e-44 |
| 178 | WMU93373.1 | N.a. | Putative metal-dependent hydrolase | TatD family hydrolase | *V. chlorellavorus* | 0 |
| 179 | WMU93374.1 | N.a. | HP | DUF6980 family protein | *V. chlorellavorus* | 3e-120 |
| 180 | WMU93375.1 | Leucine-tRNA ligase | Methionine--tRNA ligase | Methionine--tRNA ligase | *V. chlorellavorus* | 0 |
| 181 | WMU93376.1 | N.a. | HP | None | None | none |
| 182 | WMU93377.1 | N.a. | Threonylcarbamoyladenosine tRNA methylthiotransferase | tRNA (N(6)-L-threonylcarbamoyladenosine(37)-C(2))-methylthiotransferase MtaB | *V. chlorellavorus* | 0 |
| 183 | WMU93378.1 | N.a. | Nickel-responsive regulator | CopG family ribbon-helix-helix protein | *V. chlorellavorus* | 2e-29 |
| 184 | WMU93379.1 | N.a. | Nickel-responsive regulator | CopG family ribbon-helix-helix protein | *V. chlorellavorus* | 2e-32 |
| 185 | WMU93380.1 | N.a. | HP | 2 HPs | *Vampirovibrio* spp. | ≤ 4e-151 |
| 186 | WMU93381.1 | PurB Adenylosuccinate lyase | Adenylosuccinate lyase | Adenylosuccinate lyase | *V. chlorellavorus* | 0 |
| 187 | WMU93382.1 | N.a. | HP | Mechanosensitive ion channel family protein | *V. chlorellavorus* | 0 |

| | | | | | | |
|---|---|---|---|---|---|---|
| 188 | WMU93383.1 | N.a. | Glycogen operon protein | Glycogen debranching protein GlgX | *V. chlorellavorus* | 0 |
| 189 | WMU93384.1 | N.a. | HP | MerR family transcriptional regulator | *V. chlorellavorus* | 1e-124 |
| 190 | WMU93385.1 | Cell division protease | ATP-dependent zinc metalloprotease | ATP-dependent zinc metalloprotease FtsH | *V. chlorellavorus* | 0 |
| 191 | WMU93386.1 | Threonine-tRNA ligase | Proline--tRNA ligase | Proline--tRNA ligase | *V. chlorellavorus* | 0 |
| 192 | WMU93387.1 | FusA Translation elongation factors (GTPases) | Elongation factor 4 | Translation elongation factor 4 | *V. chlorellavorus* | 0 |
| 193 | WMU93388.1 | N.a. | HP | HP | *Vampirovibrio* sp. | 2e-28 |
| 194 | WMU93389.1 | N.a. | HP | 2 HPs | *Vampirovibrio* spp. | ≤ 2e-38 |
| 195 | WMU93390.1 | Peptidase M15 | HP | D-Ala-D-Ala carboxypeptidase family metallohydrolase | *V. chlorellavorus* | 9e-82 |
| 196 | WMU93391.1 | N.a. | HP | 2 HPs | *Vampirovibrio* spp. | ≤ 1e-41 |
| 197 | WMU93392.1 | N.a. | HP | 2 HPs | *Vampirovibrio* spp. | ≤ 8e-53 |
| 198 | WMU93393.1 | N.a. | HP | Phage tail protein | *Pseudomonas* sp. NFACC36 | 3e-10 |
| 199 | WMU93394.1 | N.a. | HP | 17 HPs | Multiple species | <6e-04 |
| 200 | WMU93395.1 | N.a. | HP | Phage adaptor protein | *V. chlorellavorus* | 5e-134 |
| 201 | WMU93396.1 | N.a. | HP | 2 HPs | *Vampirovibrio* spp. | ≤ 1e-31 |
| 202 | WMU93397.1 | Capsid protein | HP | P22 phage major capsid protein family protein | *V. chlorellavorus* | 0 |
| 203 | WMU93398.1 | N.a. | HP | 2 HPs | *Vampirovibrio* spp. | ≤ 6e-74 |
| 204 | WMU93399.1 | N.a. | HP | 2 HPs | *Vampirovibrio* spp. | ≤ 1e-45 |
| 205 | WMU93400.1 | Portal protein | HP | Portal protein | *V. chlorellavorus* | 0 |
| 206 | WMU93401.1 | N.a. | HP | 3 HPs | *Vampirovibrio* spp. | ≤ 4e-19 |
| 207 | WMU93402.1 | N.a. | Potassium transporter | APC family permease | *V. chlorellavorus* | 0 |
| 208 | WMU93403.1 | Phosphoglucomutase | Phosphoglucomutase | Phosphoglucomutase/phosphomannomutase family protein | *Leptolyngbya* sp. FACHB-261 | 3e-103 |
| 209 | WMU93404.1 | N.a. | K(+)-insensitive pyrophosphate-energized proton pump | Sodium-translocating pyrophosphatase | *V. chlorellavorus* | 0 |
| 210 | WMU93405.1 | N.a. | Nickel-responsive regulator | CopG family ribbon-helix-helix protein | *V. chlorellavorus* | 2e-25 |
| 211 | WMU93406.1 | N.a. | HP | Flagellar brake protein | *V. chlorellavorus* | 2e-112 |
| 212 | WMU93407.1 | RNA polymerase sigma-W factor | ECF RNA polymerase sigma factor | Sigma-70 family RNA polymerase sigma factor | *V. chlorellavorus* | 7e-145 |
| 213 | WMU93408.1 | N.a. | HP | 2 HPs | *Vampirovibrio* spp. | ≤ 4e-96 |
| 214 | WMU93409.1 | N.a. | HP | Spy/CpxP family protein refolding chaperone | *V. chlorellavorus* | 3e-89 |
| 215 | WMU93410.1 | LysU Lysyl-tRNA synthetase (class II) | Aspartate--tRNA ligase | Aspartate--tRNA ligase | *V. chlorellavorus* | 0 |
| 216 | WMU93411.1 | BaeS Signal transduction histidine kinase | Adaptive-response sensory-kinase | GAF domain-containing protein | *V. chlorellavorus* | 0 |
| 217 | WMU93412.1 | N.a. | Dephospho-CoA kinase | Dephospho-CoA kinase | *V. chlorellavorus* | 8e-141 |
| 218 | WMU93413.1 | N.a. | HP | S-layer homology domain-containing protein | *Vallitalea pronyensis* | 7e-04 |

| | | | | | | |
|---|---|---|---|---|---|---|
| 219 | WMU93414.1 | N.a. | Ribonuclease HIII | Ribonuclease HIII | *V. chlorellavorus* | 0 |
| 220 | WMU93415.1 | N.a. | HP | DUF2730 domain-containing protein | *Myxococcus* sp. XM-1-1-1 | 2e-15 |
| 221 | WMU93416.1 | N.a. | HP | 2 HPs | *Vampirovibrio* spp. | ≤ 4e-39 |
| 222 | WMU93417.1 | N.a. | Glutamyl-tRNA(Gln) amidotransferase subunit A | Asp-tRNA(Asn)/Glu-tRNA(Gln) amidotransferase subunit GatA | *V. chlorellavorus* | 0 |
| 223 | WMU93418.1 | N.a. | Bifunctional homocysteine S-methyltransferase/5 | Methylenetetrahydrofolate reductase | *V. chlorellavorus* | 0 |
| 224 | WMU93419.1 | N.a. | HP | 2 HPs | *Vampirovibrio* spp. | ≤ 4e-75 |
| 225 | WMU93420.1 | N.a. | HP | DUF5522 domain-containing protein | *V. chlorellavorus* | 2e-37 |
| 226 | WMU93421.1 | Amidophosphoribosyltransferase | Amidophosphoribosyltransferase | Amidophosphoribosyltransferase | *V. chlorellavorus* | 0 |
| 227 | WMU93422.1 | Putative phosphoribosyl formylglycinamidine (FGAM) synthase II | Phosphoribosylformylglycinamidine synthase subunit | Phosphoribosylformylglycinamidine synthase subunit PurL | *V. chlorellavorus* | 0 |
| 228 | WMU93423.1 | N.a. | HP | Bax inhibitor-1 family protein | *V. chlorellavorus* | 1e-121 |
| 229 | WMU93424.1 | Pentapeptide repeat family protein | HP | Pentapeptide repeat-containing protein | *V. chlorellavorus* | 6e-139 |
| 230 | WMU93425.1 | N.a. | HP | HP | *V. chlorellavorus* | 2e-26 |
| 231 | WMU93426.1 | SpeB Arginase/agmatinase/ formimionoglutamate hydrolase, arginase family | Agmatinase | Agmatinase | *V. chlorellavorus* | 0 |
| 232 | WMU93427.1 | SpeE Spermidine synthase | Polyamine aminopropyltransferase | Polyamine aminopropyltransferase | *V. chlorellavorus* | 0 |
| 233 | WMU93428.1 | SpeD S-adenosylmethionine decarboxylase | S-adenosylmethionine decarboxylase proenzyme | S-adenosylmethionine decarboxylase | *V. chlorellavorus* | 3e-97 |
| 234 | WMU93429.1 | N.a. | S-adenosylmethionine decarboxylase proenzyme | Adenosylmethionine decarboxylase | *V. chlorellavorus* | 3e-113 |
| 235 | WMU93430.1 | Bactoprenol glucosyl transferase | Dodecaprenyl-phosphate galacturonate synthase | Glycosyltransferase family 2 protein | *V. chlorellavorus* | 4e-172 |
| 236 | WMU93431.1 | N.a. | HP | Phospholipid carrier-dependent glycosyltransferase | *Kordia* sp. | 3e-17 |
| 237 | WMU93432.1 | N.a. | Guanine deaminase | Guanine deaminase | *V. chlorellavorus* | 0 |
| 238 | WMU93433.1 | N.a. | HP | 2 HPs | *V. chlorellavorus* | ≤ 2e-52 |
| 239 | WMU93434.1 | ATP-dependent protease gp262 | HP | MgtC/SapB family protein | *V. chlorellavorus* | 1e-173 |
| 240 | WMU93435.1 | N.a. | HP | Aminotransferase class IV | *V. chlorellavorus* | 2e-165 |
| 241 | WMU93436.1 | N.a. | HP | ATP-dependent endonuclease | *V. chlorellavorus* | 0 |
| 242 | WMU93437.1 | N.a. | HP | Ribosomal protein S18-alanine N-acetyltransferase | *V. chlorellavorus* | 2e-136 |
| 243 | WMU93438.1 | N.a. | HP | Biotin transporter BioY | *V. chlorellavorus* | 9e-158 |
| 244 | WMU93439.1 | N.a. | 5-oxoprolinase subunit A | LamB/YcsF family protein | *V. chlorellavorus* | 0 |
| 245 | WMU93440.1 | RNA binding protein | Nucleotide-binding protein | RNase adapter RapZ | *V. chlorellavorus* | 0 |
| 246 | WMU93441.1 | N.a. | Gluconeogenesis factor | Gluconeogenesis factor YvcK family protein | *V. chlorellavorus* | 0 |
| 247 | WMU93442.1 | N.a. | 3-methyl-2-oxobutanoate hydroxymethyltransferase | 3-methyl-2-oxobutanoate hydroxymethyltransferase | *V. chlorellavorus* | 0 |

| | | | | | | |
|---|---|---|---|---|---|---|
| 248 | WMU93443.1 | N.a. | HP | HP | *V. chlorellavorus* | 2e-127 |
| 249 | WMU93444.1 | N.a. | Pantothenate synthetase | Pantoate--beta-alanine ligase | *V. chlorellavorus* | 0 |
| 250 | WMU93445.1 | N.a. | HP | 2 HPs | *V. chlorellavorus* | ≤ 3e-28 |
| 251 | WMU93446.1 | MerR family transcriptional regulator | HP | MerR family transcriptional regulator | *V. chlorellavorus* | 6e-114 |
| 252 | WMU93447.1 | Iron-sulfur cluster assembly protein | FeS cluster assembly protein | Fe-S cluster assembly protein SufB | *V. chlorellavorus* | 0 |
| 253 | WMU93448.1 | N.a. | Putative ATP-dependent transporter | Fe-S cluster assembly ATPase SufC | *V. chlorellavorus* | 0 |
| 254 | WMU93449.1 | N.a. | FeS cluster assembly protein | Fe-S cluster assembly protein SufD | *V. chlorellavorus* | 0 |
| 255 | WMU93450.1 | N.a. | Cysteine desulfurase | Cysteine desulfurase | *V. chlorellavorus* | 0 |
| 256 | WMU93451.1 | N.a. | Iron-sulfur cluster assembly scaffold protein | Fe-S cluster assembly sulfur transfer protein SufU | *V. chlorellavorus* | 1e-102 |

[a]Twelve ORFs annotated as 'unnamed protein product', 'pentapeptide', 'protein of unknown function', 'hypothetical protein' and 'gp' are listed as N.a.

[b]The top functionally annotated hit is listed. When no functionally annotated homologs were detected, the number of significant ($E < 0.001$) HP hits is shown instead.

Abbreviations: ORF, open reading frame; N.a., not annotated; HP, hypothetical protein.

**Table S2**. InterPro families and Foldseek top hits of PhiMa05 ribosomal proteins.

| ORF | GenBank accession | InterPro family name | InterPro family ID | pTM/ eTM[a] | Foldseek top hit | PDB ID | Organism | Foldseek *E*-value | Foldseek Prob. |
|---|---|---|---|---|---|---|---|---|---|
| 12 | WMU93207.1 | Small ribosomal subunit protein bS1-like | IPR050437 | 0.85 | 30S ribosomal protein S1 | 7A05-A | *Vibrio vulnificus* | 3.25e-16 | 1 |
| 13 | WMU93208.1 | Large ribosomal subunit protein bL21 | IPR001787 | 0.84 | 50S ribosomal protein L21 | 7NHN-U | *Listeria monocytogenes* EGD-e | 4.11e-11 | 1 |
| 14 | WMU93209.1 | Large ribosomal subunit protein bL27 | IPR001684 | 0.81 | 50S ribosomal protein L27 | 8BUU-W | *Bacillus subtilis* subsp. *subtilis* str. 168 | 1.60e-10 | 1 |
| 35 | WMU93230.1 | Large ribosomal subunit protein bL33 | IPR001705 | 0.59 | 50S ribosomal protein L33 | 8CVM-0 | *Cutibacterium acnes* | 2.11e-4 | 1 |
| 38 | WMU93233.1 | Ribosomal protein uL11 | IPR000911 | 0.87 | 50S ribosomal protein L11 | 5O61-J | *Mycolicibacterium smegmatis* MC2 155 | 1.15e-16 | 1 |
| 39 | WMU93234.1 | Large ribosomal subunit protein uL1, bacterial-type | IPR005878 | 0.91 | 50S ribosomal protein L1 | 7P7T-F | *Enterococcus faecalis* | 2.21e-31 | 1 |
| 108 | WMU93303.1 | GTP-binding protein, ribosome biogenesis, YsxC | IPR019987 | 0.86 | GTP-binding protein YsxC | 1SUL-B | *B. subtilis* | 1.89e-22 | 1 |
| 242 | WMU93437.1 | N-acetyltransferase RimI/Ard1 | IPR006464 | 0.89 | Ribosomal-protein-alanine acetyltransferase | 5ISV-B | *Escherichia coli* O157:H7 | 1.02e-13 | 1 |

[a]AlphaFold pTM is shown for all ORFs except ORF 12 for which D-I-TASSER eTM is shown.

Abbreviations: ORF, open reading frame; pTM, predicted template modelling score; eTM, estimated template modelling score; Prob., Foldseek probability of true-positive match.
